# A computational systems biology approach identifies SLUG as a mediator of partial Epithelial-Mesenchymal Transition (EMT)

**DOI:** 10.1101/2020.09.03.278085

**Authors:** Ayalur Raghu Subbalakshmi, Sarthak Sahoo, Kuheli Biswas, Mohit Kumar Jolly

**Affiliations:** Centre for BioSystems Science and Engineering, Indian Institute of Science, Bangalore, India; Department of Physical Sciences, Indian Institute of Science Education and Research, Kolkata, India

**Keywords:** Hybrid epithelial/mesenchymal, SLUG, Cancer Systems Biology, Epithelial-Mesenchymal Transition, Mathematical modelling, Phenotypic Stability Factor

## Abstract

Epithelial-mesenchymal plasticity comprises of reversible transitions among epithelial, hybrid epithelial/mesenchymal (E/M) and mesenchymal phenotypes, and underlies various aspects of aggressive tumor progression such as metastasis, therapy resistance and immune evasion. The process of cells attaining one or more hybrid E/M phenotypes is termed as partial EMT. Cells in hybrid E/M phenotype(s) can be more aggressive than those in either fully epithelial or mesenchymal state. Thus, identifying regulators of hybrid E/M phenotypes is essential to decipher the rheostats of phenotypic plasticity and consequent accelerators of metastasis. Here, using a computational systems biology approach, we demonstrate that SLUG (SNAIL2) – an EMT-inducing transcription factor – can inhibit cells from undergoing a complete EMT and thus stabilizing them in hybrid E/M phenotype(s). It expands the parametric range enabling the existence of a hybrid E/M phenotype, thereby behaving as a phenotypic stability factor (PSF). Our simulations suggest that this specific property of SLUG emerges from the topology of the regulatory network it forms with other key regulators of epithelial-mesenchymal plasticity. Clinical data suggests that SLUG associates with worse patient prognosis across multiple carcinomas. Together, our results indicate that SLUG can stabilize hybrid E/M phenotype(s).

## Introduction

Cancer metastasis remains the major cause of cancer related deaths [Gupta, and Massagué, 2006]. Metastasis is a dynamic process with extremely high (>99.5%) attrition rates; the minority of cells successful in establishing metastases are highly plastic, i.e. capable of adapting to their environmental changes [Jolly et al., 2017b; Celià-Terrassa, and Kang, 2016]. An important axis of plasticity involved in metastasis is epithelial-mesenchymal plasticity, i.e. bidirectional transitions among epithelial, mesenchymal and one or more hybrid epithelial/ mesenchymal (E/M) phenotypes, i.e. a complete or partial epithelial mesenchymal transition (EMT) and its reverse phenomena mesenchymal epithelial transition (MET) [Jolly, and Celia-Terrassa, 2019]. Recent *in vitro*, *in vivo* and *in silico* studies suggest that EMT or MET are not ‘all-or-none’ processes; instead, cells can attain one or more hybrid E/M phenotypes which may be more plastic as compared to ‘fully’ epithelial or mesenchymal ones [Pastushenko et al., 2018; Font-Clos et al., 2018; Tripathi et al., 2020]. Consistently, cells acquiring hybrid E/M phenotype(s) are crucial for tumorigenicity and/or metastasis-initiating properties across many cancer types [Kröger et al., 2019; Pastushenko et al., 2018], and undergoing a ‘full’ EMT may lead to loss of such ‘stemness’ traits [Bierie et al., 2017; Celià-Terrassa et al., 2012]. Thus, hybrid E/M cells are likely to be the ‘fittest’ for metastasis [Jolly et al., 2018], a hypothesis that is strengthened by increasing reports suggesting an association of hybrid E/M phenotype with worse clinicopathological features [Godin et al., 2020; Liao, and Yang, 2020]. Therefore, it is critical to understand the molecular players regulating hybrid E/M phenotypes to decode their effect on aggravated metastasis and identify therapeutic approaches to restrict them.

Investigation of EMT/MET in developmental contexts have helped identify the modulators of cancer cell plasticity [Nieto et al., 2016]. One of the key molecules that has been associated with hybrid E/M phenotypes in various developmental contexts is SLUG (SNAIL2) [Jolly et al., 2015]; resonating observations have been seen in clinical settings too recently [Steinbichler et al., 2020]. However, this association has been mostly at a phenomenological level. SLUG belongs to a family of zinc-finger transcription factors SNAI. In mammals, this family consists of SNAIL (encoded by *SNAI1* gene; also called as SNAIL1), SLUG (encoded by *SNAI2* gene; also called as SNAIL2) and SMUC (encoded by *SNAI3* gene; also called as SNAIL3) [Barrallo-Gimeno, and Nieto, 2005]. While SMUC is much less-studied, SNAIL and SLUG are observed across species from *Drosophila melanogaster* to humans with functions varying from development to oncogenesis [Wu et al., 2005]. In Drosophila, they are involved in formation of mesoderm and development of the central nervous system [Hemavathy et al., 2000]. In vertebrates, they are involved in the formation of the neural crest [Hu et al., 2014], EMT of mesodermal cells and can also participate in EMT-independent functions such as preventing apoptosis from gamma-irradiation in hematopoietic cells [Barrallo-Gimeno, and Nieto, 2005]. The role of SLUG and SNAIL – the two more well-studied members of the SNAI family – has been found to be partially redundant [Hemavathy et al., 2000; Stemmler et al., 2019]. But a detailed comparison between them, at least in terms of their ability to induce a partial EMT, i.e. hybrid E/M phenotype(s), remains to be done to acquire mechanistic insights into their role in mediating the dynamics of EMP.

Here, we investigate the differences between the EMT-inducing potential of SLUG and SNAIL and investigate the contribution of SLUG in being able to maintain a hybrid E/M phenotype. Previous mathematical models developed for EMP signaling have often not distinguished between SNAIL and SLUG, and taken them as a single node [Tian et al., 2013; Hong et al., 2015; Mooney et al., 2016; Lu et al., 2013]. With increasing knowledge about the different roles of SNAIL and SLUG in influencing EMP in different contexts, including their effect on one another [Ganesan et al., 2015; Varankar et al., 2019; Nakamura et al., 2018; Gras et al., 2014; Sundararajan et al., 2019; Hamidi et al., 2020; Swain et al., 2019; Chen, and Gridley, 2013], it becomes imperative to investigate their differential roles from a detailed mechanistic perspective. Therefore, we have adopted a mechanism-based computational systems biology approach to elucidate these differences. Our results suggest that SLUG is a weaker inducer of EMT as compared to SNAIL; thus, SLUG, but not SNAIL, can often stabilize a hybrid E/M phenotype. Clinical analysis indicates high SLUG expression to be associated with worse patient survival across carcinomas, endorsing the growing evidence for a more aggressive behavior of hybrid E/M phenotype(s) .

## Results

### SNAIL is a stronger inducer of complete EMT than SLUG

First, using publicly available transcriptomics datasets (GSE80042. GSE40690, GSE43495), we quantified the extent of EMT induced by SNAIL and SLUG individually in different cell lines. We used three different methods to quantify EMT – 76GS, MLR and KS. MLR and KS score the samples on a spectrum of [0, 2] and [−1,1]; the higher the score, the more mesenchymal the sample is. On the other hand, there is no pre-defined range of scores calculated by 76GS, and the higher the 76GS score, the more epithelial the sample is [Chakraborty et al., 2020].

In LNCaP prostate cancer cells with inducible and reversible expression of SNAI1 or SNAI2 (GSE80042), those induced with SNAI1 showed a stronger extent of EMT as compared to those induced with SNAI2. This trend was consistently seen across the three scoring methods (**Fig 1A**). Similarly, in immortalized human mammary epithelial cells transfected with retroviral constructs containing SNAI1, SNAI2 and SNAI3 (HMEC-hTERT), those induced with SNAI1 showed a significantly stronger EMT induction as compared to those induced with SNAI2 or SNAI3 (GSE40690); the degree of induction was the weakest for SNAI3. Again, this trend was consistent across scoring methods (**Fig 1B**). Consistently, in human mammary epithelial cells (HMLE), transduction of SNAI1 induced a stronger EMT than that of SNAI2 or TWIST (**Fig 1C**) (GSE43495). In all these scenarios, the scores calculated across these methods correlated well with one another – while KS and MLR scores were positively correlated, 76GS scores correlated negatively with KS and with MLR scores, as expected (**Fig S1A-C**).

**Fig 1:**
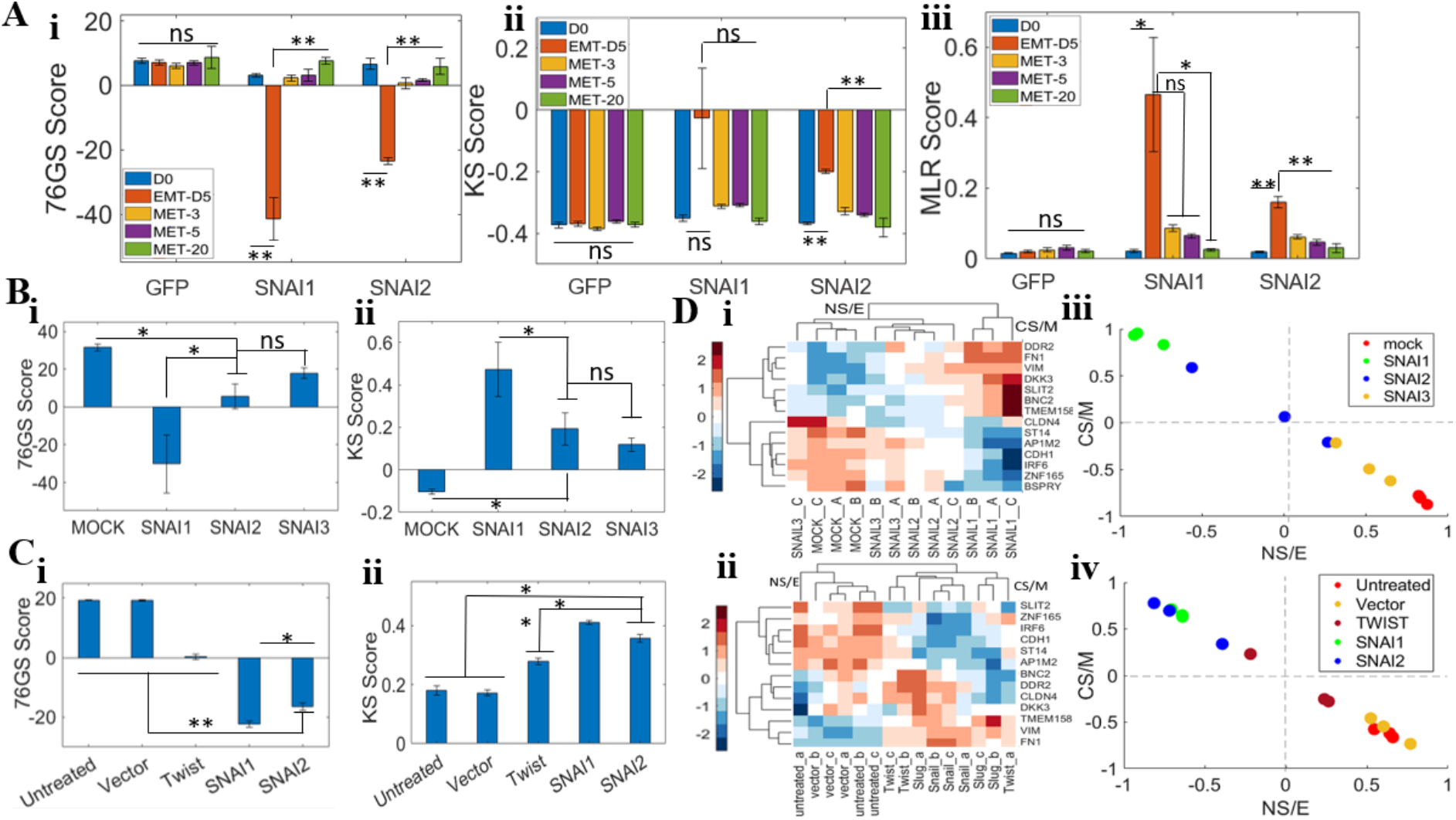
SNAIL elicits a stronger EMT than SLUG. **A)** EMT scores for GSE80042 using 76GS, KS and MLR methods **B)** EMT scores for GSE40690 using 76GS and KS methods. **C)** Sane as B) but for GSE43495. **D)** i, ii) Clustering of samples using CNCL gene list in GSE40690 and GSE43495 respectively. iii, iv) Stemness score based analysis for GSE40690 and GSE43495 respectively. * indicates a p-value < 0.05 and ** indicates p value <0.001 using the Student’s t-test with unequal variances. ‘ns’ denotes non-(statistically) significant (p >0.05).

Next, for GSE43495 and GSE40690 datasets, we also calculated the stemness scores using a gene list corresponding to EMT and stemness for breast cancer cells (CNCL) [Akbar et al., 2020]. Cells overexpressing SNAI1 were found to show a stronger enrichment of this gene signature as compared to those overexpressing SNAI2 and/or TWIST (**Fig 1D; i, ii**). When projected on the two-dimensional representation of NS/E (non-cancer-stem-like/epithelial) and CS/M (cancer stem-cell-like/mesenchymal) axes, the cells induced with SNAI1 were found to be more of CS/M^Hi^-NS/E^Low^ as compared to those induced with SNAI3 (CS/M^Low^-NS/E^Hi^) (**Fig 1D, iii**); the SNAI2 and/or TWIST-induced ones had a weaker enrichment of CS/M^Hi^-NS/E^Low^ signatures (**Fig 1D; iii, iv**). Put together, these results suggest that SNAIL is a stronger inducer of EMT than SLUG, and indicates the possibility of SLUG being associated with a partial EMT phenotype instead of a completely mesenchymal one.

### SLUG can act as a ‘phenotypic stability factor’ for a hybrid E/M phenotype

To disentangle the different effects of SNAIL vs. SLUG on EMT, we expanded the previous EMP circuitry we simulated earlier [Lu et al., 2013], by incorporating the interconnections of SLUG with miR-200 and SNAIL (**Fig 2A**). SLUG and SNAIL can inhibit each other, albeit indirectly; also, SLUG and miR-200 have been known to form a mutually inhibitory feedback loop (**Table S2**). These inter-connections were represented by a set of coupled ordinary differential equations (ODEs) (**Section S1, Table S1**).

**Fig 2:**
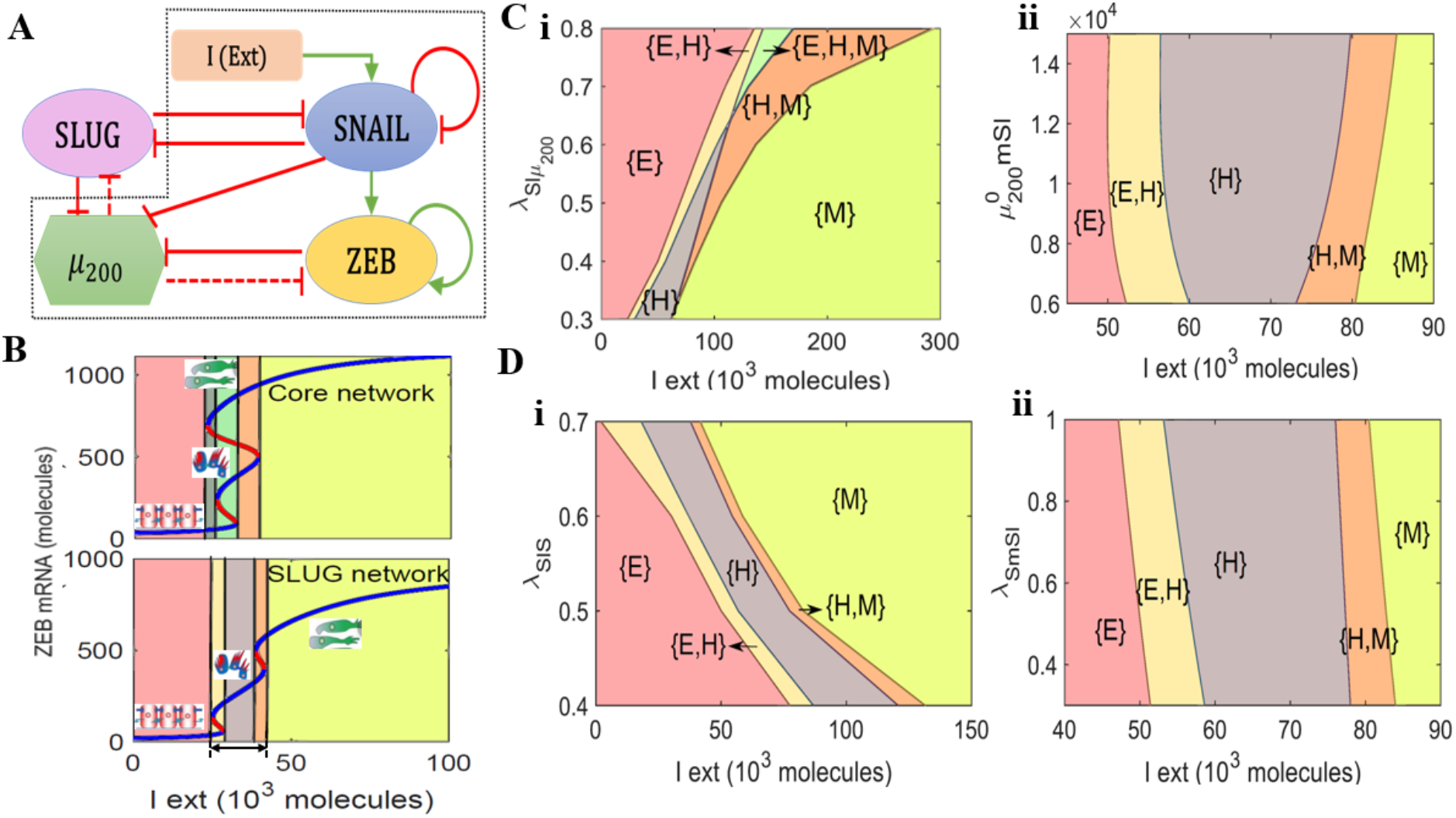
SLUG inhibits the progression to a complete EMT. **A)** Schematic representation of the EMT network coupled with SLUG. Red bars denote inhibition, green arrows indicate activation. Solid arrows represent transcriptional regulation, dash-dotted line represents indirect regulation and dotted line represent micro-RNA mediated regulation. The circuit shown within a dotted rectangle is the control case/core network (i.e. network without SLUG). **B)** Bifurcation diagrams indicating the ZEB mRNA levels for increasing external signal (I) levels for the core network (top panel) and the network including SLUG, i.e. SLUG network (bottom panel). **(C)** Phase diagrams for the SLUG network driven by an external signal (I) for varying strength of interactions along the SLUG-miR200 axis: i) Phase diagram for varying strength of repression of miR200 by SLUG, ii) Phase diagram for varying threshold level of miR200 needed for SLUG repression (i.e. reciprocal link: miR-200 inhibiting SLUG). **D)** Phase diagram for SLUG network driven by an external signal (I) for varying strength of interactions along the SLUG-SNAIL axis: i) Phase diagram for varying strength of repression of SNAIL by SLUG (ii) Phase diagram for varying strength of the reciprocal link (i.e. repression of SLUG by SNAIL).

We started with first calculating the bifurcation diagram of ZEB mRNA levels for varying levels of an external EMT-inducing signal (**Fig 2B**). As the value of EMT-inducing signal increases, the cells can switch from an epithelial state (ZEB mRNA < 100 molecules) to a hybrid E/M phenotype (100 < ZEB mRNA molecules < 600) and finally to a mesenchymal state (ZEB mRNA > 600 molecules); this trend is seen for the circuit that does not include SLUG (top panel, **Fig 2B**) as well as the one that includes SLUG (bottom panel, **Fig 2B**). However, in presence of SLUG, an interesting feature was seen which was not present in the control case – a parameter region in which the only phenotype available to cells was hybrid E/M one (grey region in bottom panel, **Fig 2B**); thus, SLUG enabled the existence of a monostable hybrid E/M phenotype. Also, in the presence of SLUG, a higher level of EMT-inducing external signal I is required to push the cells to a completely mesenchymal state, representing the effect of SLUG in preventing cells from undergoing a full EMT.

To ascertain the robustness of this prediction, we performed sensitivity analysis where each kinetic parameter of the model was varied – one at a time – by ± 10%, and the corresponding change in the range of values of the external EMT-driving signal (I) enabling a hybrid E/M phenotype (i.e. the interval of x-axis between double-sided arrows shown at the bottom) was measured (**Fig S2A**). Except a few parameter cases, the change in this interval of values of I was found to be less than 10% for a corresponding 10% change in the parameter values, indicating the robustness of the system. A change in few selected parameters such as those controlling ZEB self-activation or the effect of SLUG inhibiting SNAIL exhibited stronger sensitivity; but even in these cases, the decrease in the range of values of I for which a hybrid E/M existed was smaller as compared to the case in the absence of SLUG (dotted line in **Fig S2A**). Thus, SLUG can be thought of as a potential ‘phenotypic stability factor’ for hybrid E/M phenotype.

We next sought to examine how this trait of enabling hybrid E/M phenotype changed when the strengths of interactions between SLUG-miR200 and SLUG-SNAIL were varied. While in the presence of SLUG, the hybrid state could exist in a monostable regime ({H}), it also co-existed with epithelial and mesenchymal states, thus giving rise to the multi-stable phases (i.e. combinations of co-existing phenotypes) {E,H}, {H,M} for varying strengths of external signal. On the other hand, neither the {E, H, M} phase nor the {E, M} one was seen in the circuit containing SLUG, as opposed to the control case (compare the top vs. bottom panels in **Fig 2B**). First, we analysed the effect of interaction between SLUG and miR200 on the dynamics (**Fig 2C**). When the strength of repression of miR-200 by SLUG is reduced, two key changes are seen: a) the {M} region expands while the {E} region shrinks, i.e. it is possible for cells to exit an epithelial phenotype and/or undergo a complete EMT at lower values of external signal I, and b) the monostable hybrid region ({H}) disappears and the {E, H, M} phase emerges (**Fig 2C, i**). Similar trends, albeit to a weaker extent, were also seen when the threshold level of SLUG needed to repress miR200 is altered (**Fig S2B**). Further, to determine the effect of the reciprocal link (i.e. repression of SLUG by miR200), we varied the threshold levels of miR200 needed to repress SLUG. As this threshold value is increased, i.e. as the inhibition of SLUG by miR-200 is weakened, a larger {H} region is observed at the expense of shrinking {E} and {M} regions (**Fig 2C, ii**). Put together, these results suggests that as the influence of SLUG is reduced (i.e. weaker repression of miR-200 by SLUG, or stronger inhibition of SLUG by miR-200), the existence of a monostable hybrid state ({H}) is disfavoured. Conversely, when the effective impact of SLUG in modulating the EMP dynamics is increased (i.e. a stronger repression of miR-200 by SLUG, or weaker inhibition of SLUG by miR-200), the existence of a monostable {H} phase is facilitated.

Next, we quantified the effect of interactions between SLUG and SNAIL in altering the EMP dynamics (**Fig 2D**). As the strength of repression of SNAIL by SLUG is decreased, the {E} phase shrinks and the {M} phase expands, i.e. when SNAIL is inhibited weakly by SLUG, a milder activation of SNAIL by the external signal I is sufficient to induce an EMT (**Fig 2D, i**). However, no changes were observed when the threshold value of SLUG needed to repress SNAIL was altered (**Fig S2C**). To determine the effects of the reciprocal link (i.e. transcriptional inhibition of SLUG by SNAIL), we varied the strength of repression of the corresponding link, but did not observe any appreciable change in the phase boundaries (**Fig 2D, ii; Fig S2D**).

SLUG is also known to self-activate in certain contexts [Kumar et al., 2015; Zhou et al., 2019]. Thus, we examined the changes in EMP dynamics upon including SLUG self-activation. We observed that irrespective of the link whose strength was being varied, including self-activation of SLUG increases the parameter region for the existence of monostable {H} region. For instance, the {H} phase was seen to persist even when SLUG repressed miR-200 weakly and consequently the emergence of the tri-stable {E,H,M} phase was not noticed in presence of SLUG self-activation (compare **Fig S2E** with **Fig 2C, i**). Similarly, transition into a completely mesenchymal state ({M}) required larger values of external signal I activating SNAIL (compare **Fig S2F** with **Fig 2D, i**), leading to an expanded monostable hybrid ({H}) region. Therefore, SLUG self-activation may further enhance the likelihood of cells stably acquiring a partial EMT state and prevent them from transitioning to a fully mesenchymal state.

Finally, we replotted the abovementioned phase diagrams for a few varied values of other parameters pertaining to relative strengths of interaction of SLUG with miR-200 and/or SNAIL. Altering the degree of inhibition of SLUG by SNAIL did not significantly change the phase diagram (**Fig S3A-B**) as compared to the control case (**Fig 2D, i**). However, increasing or decreasing the strength of repression of SNAIL by SLUG modified the phase diagram. When SLUG represses SNAIL strongly, a larger value of the external signal I (that activates SNAIL) is required to induce EMT (compare **Fig S3C** with **Fig 2D, ii**). In contrast, when the repression of SNAIL by SLUG was weakened, the monostable epithelial and hybrid regions shrink, and a lower value of external signal (I) was sufficient to induce EMT (compare **Fig S3D** with **Fig 2D, ii**). Because relatively major changes in the phase diagram were observed when the inhibitory action of SLUG was altered rather than the repression on SLUG being altered, we plotted these phase diagrams for two cases: a) when SLUG inhibits both miR-200 and SNAIL strongly, b) when SLUG inhibits both miR-200 and SNAIL weakly. In the former case, the parameter region corresponding to a monostable hybrid E/M region ({H}) almost doubled; while the latter case led to a four-fold decrease in the same (compare **Fig S3E-F** with **Fig 2D,ii**). Overall, these simulations suggest that the stronger the influence of SLUG in inhibiting SNAIL and/or miR-200 levels, the higher the likelihood of cells maintaining a hybrid E/M phenotype.

### SLUG can alter the mean residence time of cells in a hybrid E/M phenotype

Previously identified ‘phenotypic stability factors’ – GRHL2, OVOL1/2, and ΔNP63α – had another dynamical trait: their presence increased the mean residence time (MRT) of cells in a hybrid E/M phenotype. MRT is the mean (or average) time spent by the cells in a particular basin of attraction – E, M and hybrid E/M here – as calculated via stochastic simulations [Biswas et al., 2019]. Thus, the larger the MRT of a phenotype, the higher the relative stability of the same. Hence, beyond enabling the parameter range of values of an external EMT-inducing signal for the existence of a hybrid E/M phenotype (as shown via bifurcation diagrams), an increase in MRT is usually thought of as a hallmark trait of a PSF. We investigated whether SLUG increases the MRT of cells in a hybrid E/M phenotype.

For the control case (i.e. circuit without SLUG), the epithelial state was found to be more stable in the {E,M} phase and the mesenchymal state is more stable in the {H,M} phase. All three states were found to be almost equally stable in the {E,H,M} phase (**Fig 3A**). Cells were seen to be capable of transitioning among these different states under the influence of noise, depending on the level of the EMT-inducing external signal (I) (**Fig 3B**). In the presence of SLUG, the hybrid state was found to be more stable than the epithelial state in the {E,H} phase and also in the {H,M} phase (**Fig 3C**), and cells switched back and forth from hybrid E/M state to epithelial or mesenchymal state depending on the value of the external signal (I) (**Fig 3D**). Thus, the presence of SLUG was capable of enabling a higher MRT for the hybrid E/M state.

**Fig 3:**
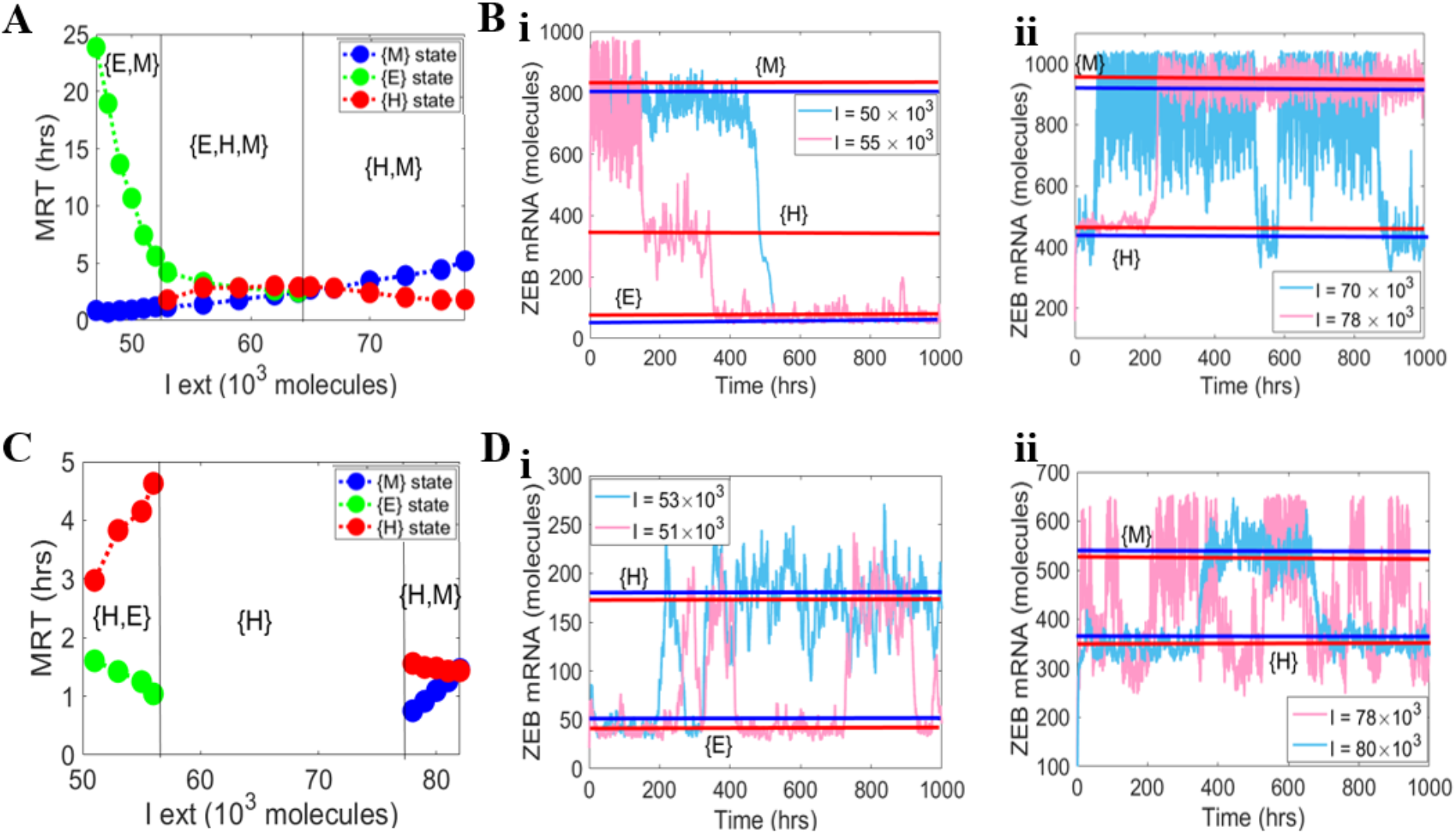
Mean residence time analysis for core network (i.e. absence of SLUG) vs. the SLUG network. **A)** Variation of mean residence time (MRT) with varying external signal (I) levels for the core network. **B)** Stochastic simulation of core circuit (I) showing switching between E and M states at I=50,000 molecules, switching among E, H and M states at I=55,000 molecules, and switching between H and M states at I=70,000 and I=78,000 molecules. The solid straight lines indicate average ZEB mRNA values for a given state and external signal strength. **C)** Same as A) but for the SLUG network **D)** Stochastic simulation for the SLUG network showing switch between E and H states at I=51,000 and I=53,000 molecules, and switching between H and M states at I=78,000 and I=80,000 molecules.

However, in the {H, M} phase, as the levels of I increase, the MRT of mesenchymal state can be larger than that of a hybrid E/M state, as the system approaches a monostable mesenchymal state ({M}). These trends are also seen in potential landscape diagrams for corresponding cases (**Fig S4**).

### The role of SLUG in stabilizing hybrid E/M state is a function of the network topology

Next, we deciphered the generic emergent properties of an extended gene regulatory network involving SNAIL, SLUG, ZEB1, miR-200 and E-cadherin (*CDH1*) **(Fig 4A)**. At a transcriptional level, ZEB1, SNAIL and SLUG can all repress E-cadherin [Batlle et al., 2000; Sterneck et al., 2020; Mooney et al., 2016], and SNAIL and SLUG can activate ZEB1 [Wels et al., 2011; Wu et al., 2017]. Further, SLUG is known to self-activate [Kumar et al., 2015], but SNAIL self-inhibits [Peiró et al., 2006]. E-cadherin can also indirectly repress ZEB1 by regulating the localization of β-catenin [Mooney et al., 2016].

**Fig 4:**
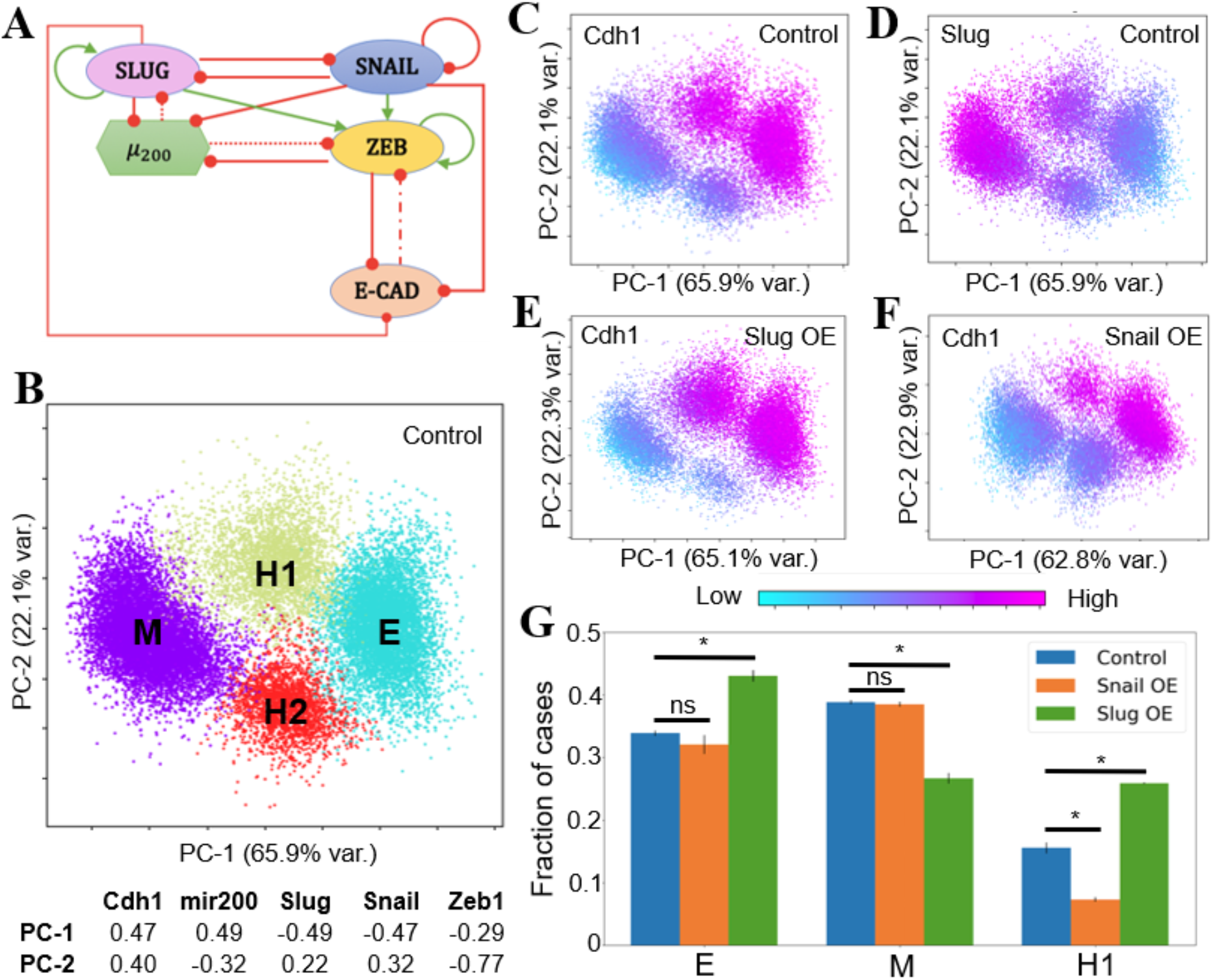
RACIPE analysis reveals the existence of a SLUG^+^CDH1^+^ hybrid E/M phenotype. **A)** Schematic of the extended gene regulatory network showing the interactions of SNAIL and SLUG with ZEB1, miR200 and CDH1. The green links represent activating links, and the red links represent inhibition links. Dotted line shows the microRNA-mediated regulation. Dashed-dotted line shows indirect inhibition of ZEB by E-cadherin through β-catenin. **B)** Principal Component Analysis (PCA) plot showing the different clusters that emerge from the RACIPE analysis, also identified via hierarchical clustering for the RACIPE output. E and M represent the epithelial and the mesenchymal phenotypes respectively. H1 and H2 are two hybrid phenotypes. Contributions of the various genes to the two principal component axes PC1 and PC2 are listed (PC1 represents the canonical EMT axis). **C)** PCA plot coloured by the expression levels of CDH1. **D)** PCA plot coloured by the expression levels of SLUG. **E)** PCA plot from RACIPE analysis in the case of SLUG over-expression (coloured by the gene expression levels of CDH1). **F)** PCA plot from RACIPE analysis in the case of SNAIL over-expression (coloured by the gene expression levels of CDH1). For E-F, the PC1 obtained is qualitatively similar to PC1 obtained for the control (panels B-D) case; also, PC1 and PC2 explain similar variance, indicating no major changes in structure of the steady state solution set. **G)** Quantification of the frequency of steady state solutions for the clusters corresponding to E, M and H1 showing that H1 phenotype is depleted by SNAIL over-expression while it is enriched for by SLUG over-expression (* indicates a p-value < 0.01 by using the Student’s t-test with unequal variances; ‘ns’ denotes non-(statistically) significant change). Error bars represent the mean +/− standard deviation for results obtained from n=3 independent RACIPE replicates.

We used a computational framework Random Circuit Perturbation Analysis (RACIPE) that enables mapping the general dynamic properties of gene regulatory networks by solving a set of coupled ordinary differential equations (ODEs) to obtain the possible steady state solutions for a given network and corresponding ensemble of parameter sets. All the parameters and initial conditions required for solving the set of ODEs are randomly sampled from biologically relevant ranges [Huang et al., 2018]. We performed RACIPE analysis on this gene regulatory network (**Fig 4A**) to find the robust steady state solutions emerging from it, for an ensemble of parameter sets. We performed Principal Component Analysis (PCA) on all the steady state solutions produced by RACIPE (**Fig 4B**). The PCA plot reveals the existence of four distinct clusters: Epithelial (E), Mesenchymal(M) and two hybrid clusters (H1 and H2), as confirmed by hierarchical clustering on all the steady state solutions obtained via RACIPE (**Fig 4B**). PC1 explains more than 65% of the variance in the data, and can be considered as the “canonical” EMT axis, given its composition that places CDH1 and miR-200 (epithelial markers) in one group and ZEB1, SLUG and SNAIL (mesenchymal markers) in the other opposing one. Thus, E and M clusters are well-segregated along the PC1 axis, and two clusters potentially representing hybrid E/M states are sandwiched between the E and M ends. These two clusters (H1, H2) seem to be segregated along PC2 axis. Biologically, the hybrid cluster H1 can denote a SLUG+ CDH1+ cell phenotype which has been documented recently experimentally as well (**Fig 4C-D**) [Sterneck et al., 2020].

Next, we compared the effect of overexpression of SLUG vs. SNAIL. Interestingly, these both scenarios showed opposite results: while SLUG overexpression enriched the SLUG+ CDH1+ phenotype, SNAIL overexpression depleted it (**Fig 4E-G**), suggesting that SNAIL and SLUG may play different or even opposing roles in driving EMP, owing to the differences in underlying network topology. Furthermore, this enrichment of the SLUG+ CDH1+ population in the case of SLUG over-expression happens at the expense of the mesenchymal (M) cell population (**Fig 4G**), lending credence to our previous results that that SLUG is associated specifically with a partial EMT instead of a full EMT phenotype. Besides, this analysis endorses that network topology can be used to pinpoint potential PSFs for the hybrid E/M phenotypes [Jolly et al., 2016].

### High SLUG expression correlates with poor prognosis

Hybrid E/M phenotype is thought of as more aggressive [Pastushenko, and Blanpain, 2019; Jolly et al., 2018]. This hypothesis is further corroborated by the data showing poor patient survival for higher expression of PSFs such as GRHL2, OVOL1/2 and NRF2 [Jolly et al., 2016; Bocci et al., 2019]. To ascertain the effects of SLUG on patient prognosis we investigated its role in patient outcomes. We observed that high SLUG expression associated consistently with poor relapse-free survival in lung, ovarian and colorectal cancer (**Fig 5A-D**), and with worse overall survival in lung, ovarian, colorectal and pancreatic cancer (**Fig 5E-H, Fig S5A-C**), indicating that high SLUG levels can correlate with poor prognosis in many carcinomas.

**Fig 5:**
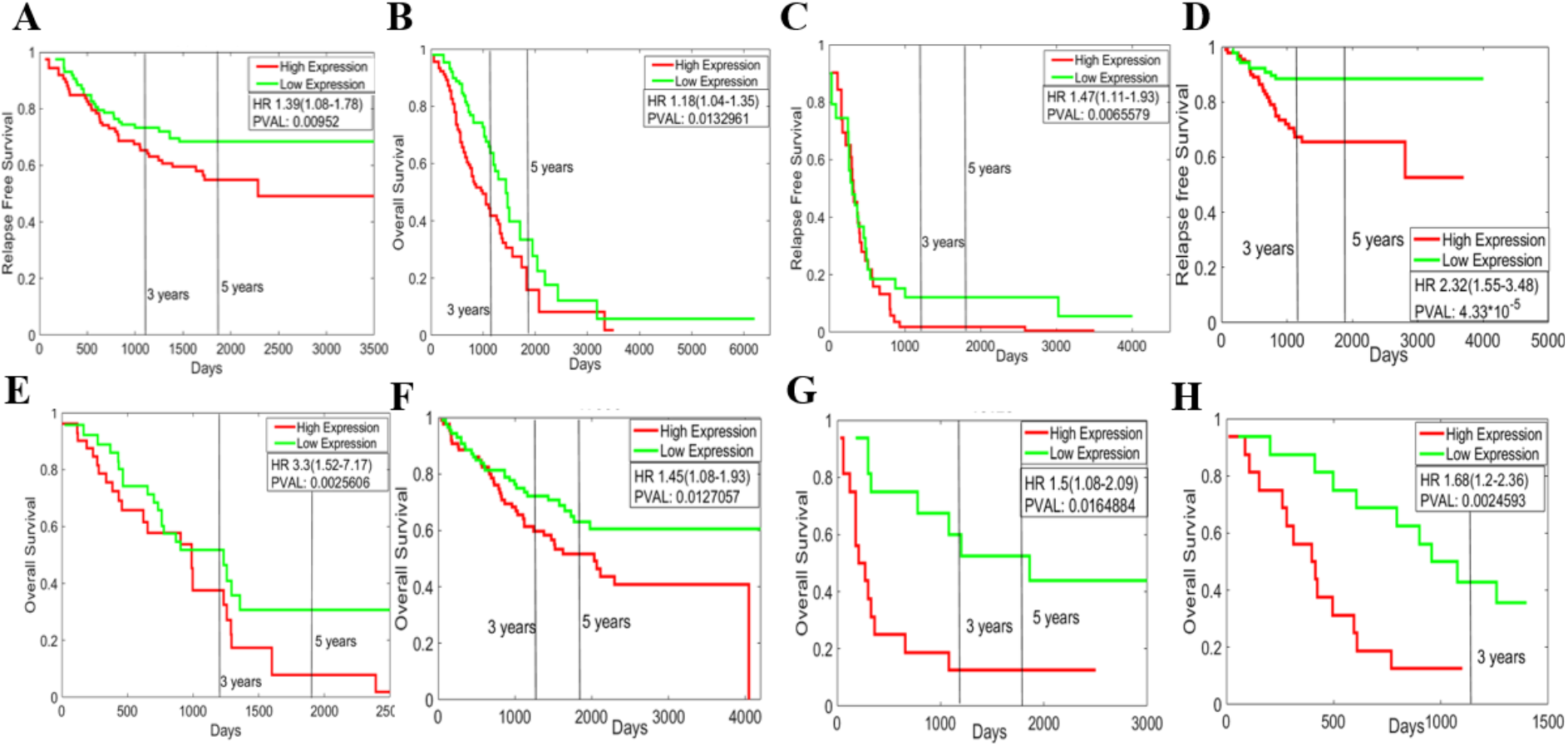
SLUG correlates with poor patient survival. **A-D)** Relapse free survival of GSE31210 (lung cancer samples), GSE9891 (ovarian cancer samples), GSE30161 (ovarian cancer sample) and GSE14333 (colorectal cancer sample) respectively. **E-H)** Overall survival of TCGA-LUAD (lung cancer sample), GSE17536 (colorectal cancer sample), GSE16125 (colorectal cancer sample) and GSE50827 (pancreatic cancer sample). HR denotes hazard ratio; PVAL denotes p-value.

In the case of breast cancer, the results were equivocal. High SLUG expression correlated with better overall patient survival (**Fig S5D**) and better metastasis-free survival (**Fig S5E,F**), but with poor relapse free survival (**Fig S5G,H**). These context-specific differences of the association of SLUG with breast cancer prognosis traits remains to be better understood.

## Discussion

Cancer cell plasticity and metastasis are complex and emergent phenomena whose molecular underpinnings are yet to be comprehensively identified. An important axis of cellular plasticity is epithelial-mesenchymal plasticity (EMP). Various families of transcription factors have been known to affect EMP – TWIST1/2, ZEB1/2, and SNAIL1/2 can induce EMT [Lamouille et al., 2014], while OVOL1/2 and GRHL1/2/3 can induce MET [Sundararajan et al., 2020; Saxena et al., 2020]. Recent investigations have highlighted different and possibly even a contradictory role of different members of the same transcription factor family, for instance, the opposing roles of ZEB1 and ZEB2 in melanoma [Caramel et al., 2013]. Here, we examined the differences in the roles of SNAIL1 (encoded by the *SNAI1* gene; also called as SNAIL) and SNAIL2 (encoded by the *SNAI2* gene; also called as SLUG) in inducing EMT.

Our results suggest that SLUG, but not necessarily SNAIL, associates with a partial EMT or hybrid E/M phenotype. These observations are reminiscent of experiments suggesting that SNAIL is a stronger inducer of molecular and/or morphological features associated with EMT, as compared to SLUG and SNAIL3 [Gras et al., 2014]. Similar to other EMT-TFs, SLUG initiates EMT by repressing *CDH1* by binding to the E-boxes [Hajra et al., 2002; Slabáková et al., 2011]; however, its binding affinity to the corresponding E-box DNA element was found to be two order of magnitude less than that of SNAIL [Bolós et al., 2003]. Intriguingly, in TGFβ1-induced EMT in BPH-1 cells, SLUG expression was found to be decreased at later time points (72-96 hours), but SNAIL and ZEB1 levels were consistently high [Slabáková et al., 2011], indicating a possible role of SLUG is initiating rather than maintaining EMT [Savagner et al., 1997]. Consistently, SLUG levels were observed to peak in intermediate states of EMT in HMLE cells treated with TGFβ [van Dijk et al., 2018], endorsing the specific association of SLUG with a partially differentiated phenotype as witnessed in multiple scenarios involving EMT – embryonic development, wound healing and carcinoma progression [Leroy, and Mostov, 2007; Savagner et al., 2005; Hudson et al., 2009; Côme et al., 2006]. SLUG is also known to activate P-cadherin [Idoux-Gillet et al., 2018], another proposed marker for partial EMT [Ribeiro, and Paredes, 2015].

The association of SLUG with partial EMT phenotype(s) also helps contextualize the growing literature of hybrid E/M phenotype being more stem-like as compared to a complete EMT [Jolly et al., 2019], and the functional role of SLUG in enabling stemness [Sterneck et al., 2020]. Mammary stem cells exhibit high SLUG levels, and SLUG-expressing normal mammary epithelial cells were seen to exhibit a partial EMT phenotype and generated 17 times more organoids than the control case [Guo et al., 2012; Ye et al., 2015]. Upon knockdown of SLUG, mammary stem cells are unable to transition to basal progenitor cells, leading to abnormal mammary architecture [Phillips et al., 2014]. The mammosphere forming potential of basal progenitor cells also decreased upon SLUG knockdown, thus the primary mammospheres generated from SLUG-KO cells were unable to give rise to secondary mammospheres, highlighting the role of SLUG on self-renewal processes in mammary gland morphogenesis [Nassour et al., 2012]. Consistent observations have been made in breast cancer and prostate cancer, where SLUG degradation promotes the differentiation of breast cancer stem cells [Fraile et al., 2020], and the basal prostate cells that expressed SLUG exhibited a partial EMT phenotype and displayed increased stemness ability [Kahonouva et al., 2020]. The functional contribution of SLUG *in vitro* and/or *in vivo* enabling stemness is also observed in human epidermal progenitor cells [Mistry et al., 2014] as well as other cancers such as glioblastoma [Chesnelong et al., 2019], hepatocellular carcinoma [Tang et al., 2016; Sun et al., 2014], colorectal cancer [Kato et al., 2020], lung cancer [Kim et al., 2020] and squamous cell carcinomas [Yu et al., 2016; Moon et al., 2020]. Whether higher stemness is connected to a hybrid E/M phenotype in these cancers remains to be identified.

Our results suggest that the network topology that SLUG forms with SNAIL and miR-200 enables its role as a ‘phenotypic stability factor’ for a hybrid E/M phenotype. It should be noted the strengths of interactions of SLUG with SNAIL and/or miR-200 can be context-dependent, thus altering its ability to be able to stabilize hybrid E/M phenotype(s). miR-200 family members have been shown to inhibit SLUG in esophageal cancer and prostate cancer [Liu et al., 2013; Zong et al., 2019; Basu et al., 2020], while SLUG can inhibit miR-200 in prostate cancer [Liu et al., 2013]. On the other hand, SLUG and SNAIL have been reported to inhibit each other in oral cancer [Nakamura et al., 2018] and breast cancer [Ganesan et al., 2015]. The effect of SLUG on EMP dynamics can also be influenced by upstream regulators of SLUG such as RNF8, YAP, C/EBPδ, miR-151, CBP, RCP, HDAC1, HDAC3, BRD4 and STAT3 [Cheng et al., 2020; Chesnelong et al., 2019; Kuang et al., 2020; Wang et al., 2020; Daugaard et al., 2017; Jury et al., 2020; Dai et al., 2020; Hwang et al., 2017; Hu et al., 2020; Shafran et al., 2020]. Moreover, SLUG may be involved in feedback loops with one or more p63 isoforms [Srivastava et al., 2018; Dang et al., 2016; Herfs et al., 2010] that can also enable a partial EMT [Jolly et al., 2017a; Westcott et al., 2020]. Thus, while the proposed role of SLUG in stabilizing hybrid E/M phenotype(s) is largely robust to parametric variations, including other links and/or nodes at can alter the landscape of EMP dynamics. A better understanding of similarities and differences in terms of networks activated by SNAIL vs. SLUG to influence the different states of EMT and associated traits such as drug resistance [Kurrey et al., 2009; Li et al., 2020], immune evasion [Tripathi et al., 2016], anoikis resistance [Huang et al., 2013] and autophagy [Xu et al., 2019] across different cancer types is therefore required.

## Materials and Methods

### Software and Datasets

All computational analysis were performed using MATLAB (version 2018b) and microarray datasets were downloaded using *GEOquery* R Bioconductor package [Davis, and Meltzer, 2007]. Pre-processing of the microarray datasets was performed for each sample to obtain the gene-wise expression from probe-wise expression matrix using R (version 4.0.0).

### EMT score calculation

The epithelial-mesenchymal transition (EMT) scores for each dataset was calculated using 76GS, KS and MLR methods as described earlier [Chakraborty et al., 2020]

#### 76GS

A weighted sum of 76 gene expression levels was used to calculate the score for each sample. The mean across all tumor samples was subtracted to center the scores so that the grand mean of the score was zero. Negative scores can be interpreted as M phenotype whereas the positive scores as E.

#### MLR

The EMT status by the MLR method is predicted based on order of the categorical structures and the principle that the hybrid E/M state falls in a region intermediary to E and M. The sample scores range between 0 to 2 where 0 corresponds to pure E and 2 corresponds to pure M, and a score of 1 indicates a maximally hybrid phenotype. These scores are calculated based on the probability of a given sample being assigned to the E, E/M, and M phenotypes.

#### KS

The scores were calculated by comparing cumulative distribution functions (CDFs) of E and M signatures. If the score is positive, then the sample is said to be mesenchymal (M) and if it is negative, it is considered epithelial (E).

### Stemness score calculation

The stemness score was calculated using CNCL as described [Akbar et al., 2020]. To identify the stem-cell-like (CS/M) and non-stem-cell like (NS/E) cells, the samples were clustered based on the expression of CSC/non-CSC gene list (CNCL). The quantitative stemness score was calculated by the correlation of CNCL gene expression values for each sample to the CS/M and NS/E matrices generated based on the median expression levels for each gene in the CNCL for CS/M and also for NS/E cells in the CCLE database. The two Pearson’s correlation values generated for each matrix, CS/M(r) and NS/E(r), are plotted.

### Mathematical Modelling

As per the schematic shown in Fig 2A, the dynamics of all four molecular species (miR-200, SNAIL, ZEB and SLUG) was described by a system of coupled ODEs. The generic chemical rate equation given below describes the level of a protein, mRNA or micro-RNA (X):

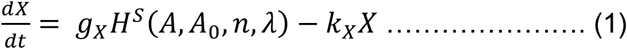

Where the first term g_x_ signifies the basal rate of production; the terms multiplied to g_x_ represent the transcriptional/translational/post-translational regulations due to interactions among the species in the system, as defined by the Hills function *H*^*S*^(*A*, *A*_0_, *n*, λ). The rate of degradation of the species (X) is defined by the term k_X_X based on first order kinetics. The complete set of equations and parameters are presented in the Supplementary Material.

### Mean Residence Time Analysis

As the degradation rate of ZEB mRNA and SLUG mRNA is much greater than that of ZEB protein and miR-200, and also the production rate of SLUG and SNAIL protein is much more larger than that of ZEB protein and miR-200, we assumed that ZEB mRNA, SNAIL protein SLUG mRNA and SLUG protein reach to the equilibrium much faster relatively, i.e. 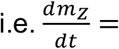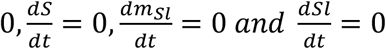. This assumption reduces the equations given in Supplementary Material to two coupled ODEs of ZEB and miR-200. Then, the dynamical system was simulated to obtain the time evolution of ZEB protein and miR-200 in presence of external noise using Euler–Maruyama simulation. Dynamical states of the system were coarse grained as an itinerary of basins visited from the time evolution of ZEB and miR-200. MRT was calculated by multiplying total number of successive states with it [Biswas et al., 2019].

### RACIPE (Random Circuit Perturbation)

We performed Random Circuit Perturbation analysis on the extended circuit (Fig4A) using the default settings for 10000 parameter sets. We considered all RACIPE steady state solutions up to tetra-stable parameter sets for PCA plots. PCA was performed on all the steady state solutions from the RACIPE output data. The clusters were identified by performing hierarchical clustering on the RACIPE data by setting the number of clusters to 4 and the number of steady states solutions ending up in each of these clusters were quantified. We also performed RACIPE on the same circuit but with either over-expressing SNAIL or SLUG by 10-fold. The same workflow as described above was used to analyze the over-expression based analysis.

### Kaplan–Meier Analysis

ProgGene [Goswami, and Nakshatri, 2014] was used for conducting Kaplan–Meier analysis. The number of samples in SLUG-high vs. SLUG-low categories are given below:

GSE31210 (Lung cancer sample): n(High) = 113, n(Low) = 113

GSE9891 (Ovarian cancer sample): n(High) = 138, n(Low) = 138

GSE30161 (Ovarian cancer sample): n(High) = 29, n(Low) = 29

GSE14333 (Colorectal cancer sample): n(High) = 94, n(Low) = 93

TCGA-LUAD (Lung cancer sample): n(High) = 75, n(Low) = 75

GSE17536 (Colorectal cancer sample): n(High) = 87, n(Low) = 87

GSE16125 (Colorectal cancer sample): n(High) = 16, n(Low) = 16

GSE50827 (Pancreatic cancer sample): n(High) = 16, n(Low) = 16

GSE26712 (Ovarian cancer sample): n(High) = 93, n(Low) = 92

GSE18520 (Ovarian cancer sample): n(High) = 27, n(Low) = 26

GSE13876 (Ovarian cancer sample) : n(High) = 207, n(Low) = 207

GSE3143 (Breast cancer sample): n(High) = 80, n(Low) = 77

GSE2034 (Breast cancer sample): n(High) = 143, n(Low) = 143

GSE19615(Breast cancer sample): n(High) = 58, n(Low) = 57

GSE11121 (Breast cancer sample): n(High) = 103, n(Low) = 97

GSE2990 (Breast cancer sample): n(High) = 51, n(Low) = 50

## Acknowledgements

This work was supported by Ramanujan Fellowship (SB/S2/RJN-049/2018) awarded to MKJ by Science and Engineering Research Board (SERB), Department of Science and Technology (DST), Government of India.

## Author contributions

MKJ designed and supervised research; ARS, SS and KB performed research and analyzed data; all authors contributed to writing and editing of the manuscript.

