## Supplementary Tables and Figures for "A computational systems biology approach identifies SLUG as a mediator of partial Epithelial-Mesenchymal Transition (EMT)"

**Supplementary Information**

### S1. Mathematical modelling

The dynamics of the SLUG circuit describes the dynamics of the molecular species of the EMT regulatory circuit (miR-200, Snail, Zeb) and SLUG as shown in Fig 2A. This set-up extends the mathematical model of EMT circuit previously developed (1). The following set of coupled ordinary differential equations (ODEs) represent the dynamics of the species of the circuit (External signal: I, miR-200: $\mu_{200}$, SNAIL: S, ZEB: Z, SLUG: Sl):

$$\frac{d\mu_{200}}{dt} = g_{\mu_{200}} H^s(Z,\lambda_{Z,\mu_{200}}) H^s(S,\lambda_{S,\mu_{200}}) H^s(Sl,\lambda_{Sl,\mu_{200}}) - m_Z Y_\mu(\mu_{200}) - m_{Sl} Y_\mu(\mu_{200}) - k_{\mu_{200}}\mu_{200}$$

$$\frac{dm_Z}{dt} = g_{m_Z} H^s(Z,\lambda_{Z,m_Z}) H^s(S,\lambda_{S,m_Z}) - m_Z Y_Z(\mu_{200}) - k_{m_Z} m_Z$$

$$\frac{dZ}{dt} = g_Z m_Z L(\mu_{200}) - k_Z Z$$

$$\frac{dS}{dt} = g_S H^s(I,\lambda_{I,S}) H^s(Sl,\lambda_{Sl,S}) H^s(S,\lambda_{S,S}) - k_S S$$

$$\frac{dm_{Sl}}{dt} = g_{m_{Sl}} H^s(S,\lambda_{S,m_{Sl}}) - m_{Sl} Y_Z(\mu_{200}) - k_{m_{Sl}} m_{Sl}$$

$$\frac{dSl}{dt} = g_{Sl} m_{Sl} L(\mu_{200}) - k_{Sl} Sl$$

where $g_X$ is the corresponding innate production rate and $k_X$ is the innate degradation rate.

$m_Z L(\mu_{200})$ is the net translation rate, $m_Z Y_m(\mu_{200})$ is the total ZEB mRNA active degradation rate and $m_Z Y_\mu(\mu_{200})$ is the total miR active degradation rate. $H^S$ is the shifted Hill function, defined as

$H^S(B,\lambda) = H^-(B)+\lambda H^+(B)$ ,

$H^-(B) = 1/[1+(B/B_0)^{nB}]$ ,

$H^+(B) = 1-H^-(B)$,

λ is the fold change from the basal synthesis rate due to protein B. λ >1 for activators, while λ<1 for inhibitors. When SLUG self-activation is included, the equations of SLUG are updated to include a shifted Hill function in the $\frac{dm_{Sl}}{dt}$ (SLUG mRNA) equation.

**Parameter Estimation:**

The model parameters were adopted from previously published literature for the molecular species of the core circuit (I, miR-200, Snail, Zeb) and SLUG interactions, as given below:

| Parameter | Value | Reference |
|---|---|---|
| $g\mu200$ (Molecules/Hour) | 2.1K | (1) |
| $gmZ$ (Molecules/Hour) | 11 | (1) |
| $Z^0\mu_{200}$ (Molecules) | 220K | (1) |
| $Z^0m_Z$ (Molecules) | 25K | (1) |
| $n_{Z,\mu_{200}}$ | 3 | (1) |
| $n_{Z,m_Z}$ | 2 | (1) |
| $n_{\mu_{200}}$ | 6 | (1) |
| $n_{S,\mu_{200}}$ | 2 | (1) |

| | | |
|---|---|---|
| $n_{S,m_Z}$ | 2 | (1) |
| $\lambda_{Z,\mu_{200}}$ | 0.1 | (1) |
| $\lambda_{Z,m_Z}$ | 7.5 | (1) |
| $\lambda_{S,\mu_{200}}$ | 0.1 | (1) |
| $\lambda_{S,m_Z}$ | 10 | (1) |
| $k_{\mu_{200}}$(Hour$^{-1}$) | 0.05 | (1) |
| $k_{m_Z}$(Hour$^{-1}$) | 0.5 | (1) |
| $k_Z$(Hour$^{-1}$) | 0.1 | (1) |
| $g_Z$(Hour$^{-1}$) | 0.1K | (1) |
| $S^0_{\mu_{200}}$(Molecules) | 180K | (1) |
| $S^0_{m_Z}$(Molecules) | 180K | (1) |
| $\mu^0_{200}$(Molecules) | 10K | (1) |
| $g_S$ | 18000 | (1) |
| $k_S$ | 0.125 | (1) |
| $g_{Sl}$ | 50000 | Estimated |
| $k_{Sl}$ | 0.1155 | (2) |
| $g_{m_{Sl}}$ | 90 | Estimated |
| $k_{m_{Sl}}$ | 0.5 | Estimated |
| $\lambda_{Sl,\mu_{200}}$ | 0.4 | (3) |
| $\lambda_{Sl,S}$ | 0.5 | (4) |
| $\lambda_{Sl,m_{Sl}}$ | 4 | (5) |
| $\lambda_{S,S}$ | 0.4 | (6) |
| $\lambda_{S,m_{Sl}}$ | 0.5 | (4) |
| $n_{Sl,\mu_{200}}$ | 1 | (3) |
| $n_{Sl,S}$ | 3 | (7) |
| $n_{Sl,m_{Sl}}$ | 4 | (5) |
| $n_{S,S}$ | 5 | (7) |
| $n_{S,m_{Sl}}$ | 1 | (7) |
| $Sl^0_{\mu_{200}}$ | 220000 | Estimated |
| $Sl^0_S$ | 225000 | Estimated |
| $Sl^0_{m_{Sl}}$ | 250000 | Estimated |
| $S^0_S$ | 180000 | Estimated |
| $S^0_{m_{Sl}}$ | 180000 | Estimated |
| nIs | 2 | (8) |
| $I^0S$ | 100000 | (8) |
| $\lambda_{I,S}$ | 3 | (8) |

**Table S1:** Parameter values used for simulations

The details of each link used in the network can be found in Table S2.

| **Interaction** | **Reference** |
|---|---|
| ZEB represses miR200 | (9) |
| miR200 represses ZEB | (9) |
| SNAIL represses itself | (6) |

| | |
|---|---|
| SNAIL activates ZEB | (10) |
| SNAIL represses E-cad | (11) |
| ZEB represses E-cad | (12) |
| E-cad inhibits β-catenin | (13) |
| β-catenin activates ZEB | (14) |
| SLUG self activates | (5) |
| SLUG and miR200 mutually inhibit each other | (3) |
| SLUG inhibits E-cad | (15) |
| SLUG and SNAIL mutually inhibit each other | (4) |
| ZEB self activates | (16) |
| SNAIL represses miR200 | (17) |

**Table S2:** References for specific nodes and links in the network

**Supplementary figures:**

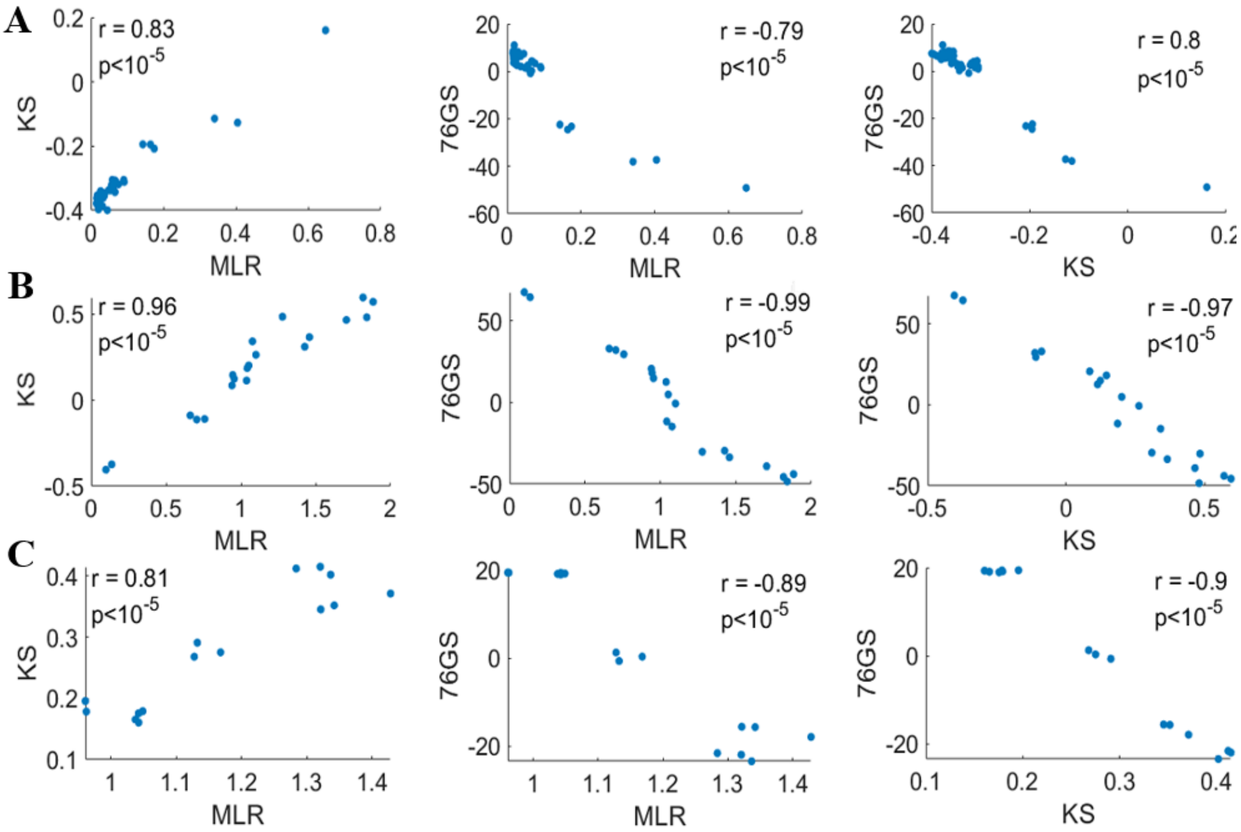


**Fig S1**: **Correlation plots among EMT scoring methods – KS, 76GS, MLR**. **A)** Spearman's correlation plots between KS, MLR and 76GS methods for GSE80042. **B)** Same as A) but for GSE40690. **C)** Same as A) but for GSE43495. Correlation coefficient is denoted by r, p-value is denoted by p.

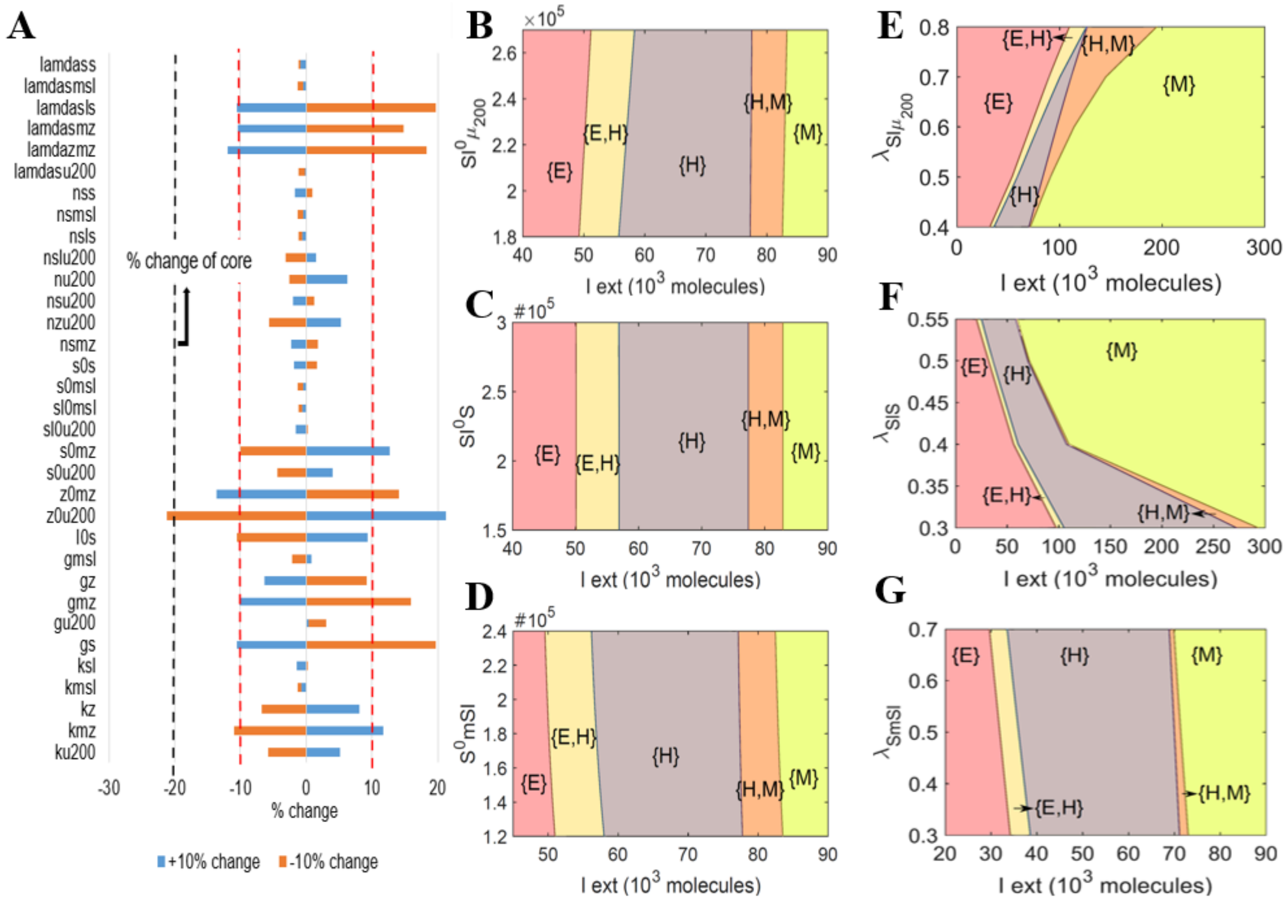


**Fig S2**: **Sensitivity analysis. A)** Sensitivity analysis indicating percentage change in the interval of external signal levels for the existence of stable hybrid E/M region, when corresponding parameter values are varied by ±10%. Black dotted line indicates the percent change in the stable hybrid region in the absence of SLUG (core network) when compared to the network that includes SLUG. **B)** Phase diagram of network including SLUG, when driven by external signal (I) and varying threshold of SLUG level for miR200 inhibition. **C)** Phase diagram of the SLUG network when driven by external signal (I) and varying threshold of SLUG level for SNAIL inhibition. **D)** Phase diagram of the SLUG network when driven by external signal (I) and varying threshold of SNAIL levels for SLUG mRNA inhibition. **E)** Same as Fig 2C, i but in presence of SLUG self-activation. **F)** Same as Fig 2D, i but in the presence of self-activation. **G)** Same as Fig 2D, ii but in the presence of SLUG self-activation. Parameters for SLUG self-activation are given in SI Table 1.

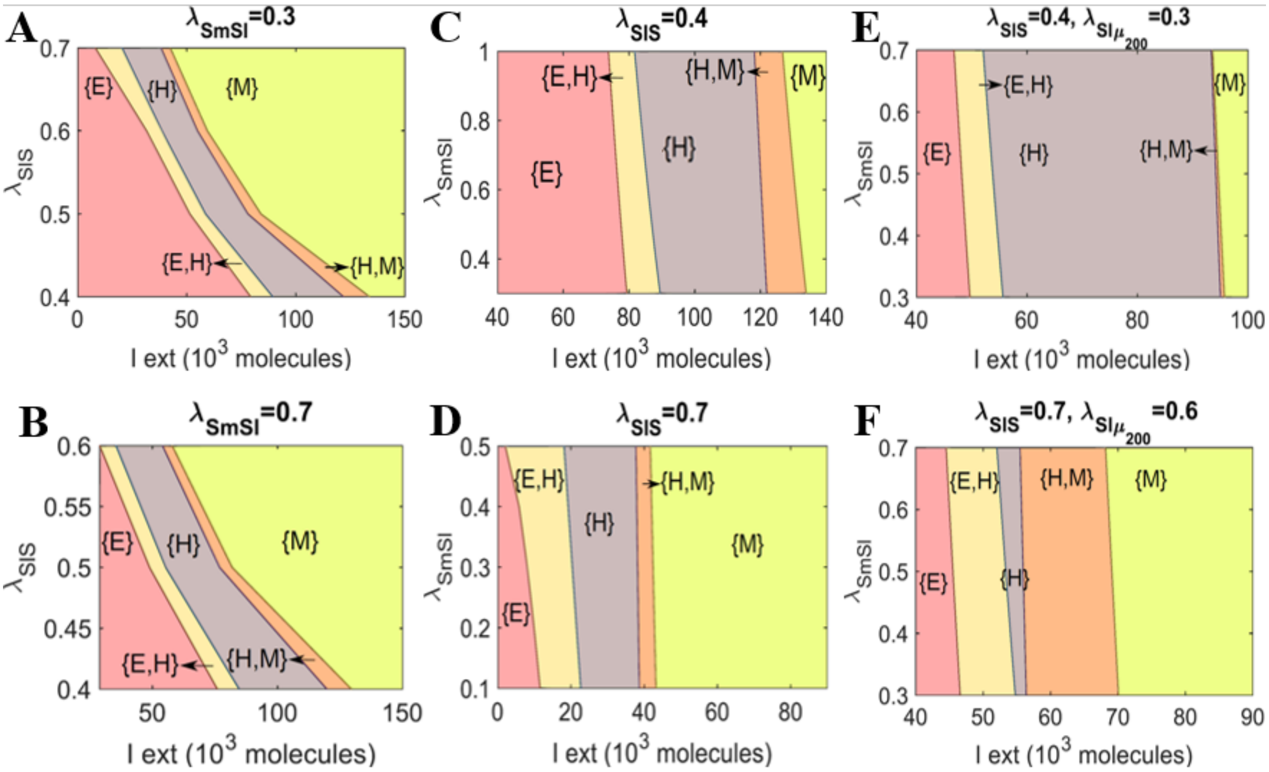


**Fig S3**: **Phase diagrams of the network including SLUG**. **A)** Phase diagram of SLUG network when driven by external signal (I) and varying strength of interaction between SLUG and SNAIL, under the conditions of a stronger inhibition of SLUG by SNAIL ( $\lambda_{S,m_{Sl}}$ = 0.3 instead of 0.5 as in Fig 2D,i). **B)** Same as A) but for $\lambda_{S,m_{Sl}}$=0.7. **C)** Phase diagram of SLUG network when driven by external signal (I) and varying strength of interaction between SLUG and SNAIL, under the conditions of a stronger inhibition of SNAIL by SLUG ( $\lambda_{Sl,S}$ = 0.4 instead of 0.5 as in Fig 2D, ii). **D)** Same as C) but for $\lambda_{Sl,S}$ = 0.7. **E)** Phase diagram of SLUG network when driven by external signal (I) and varying strength of interaction between SLUG and SNAIL, under conditions of a stronger inhibition of SNAIL and miR200 by SLUG ( $\lambda_{Sl,S}$ = 0.4 and $\lambda_{Sl,\mu_{200}}$=0.3 instead of 0.5 and 0.4 respectively as in Fig 2D, i). **F)** Same as E) but for weaker inhibition of SNAIL and miR-200 by SLUG $\lambda_{Sl,S}$ = 0.7 and $\lambda_{Sl,\mu_{200}}$=0.6.

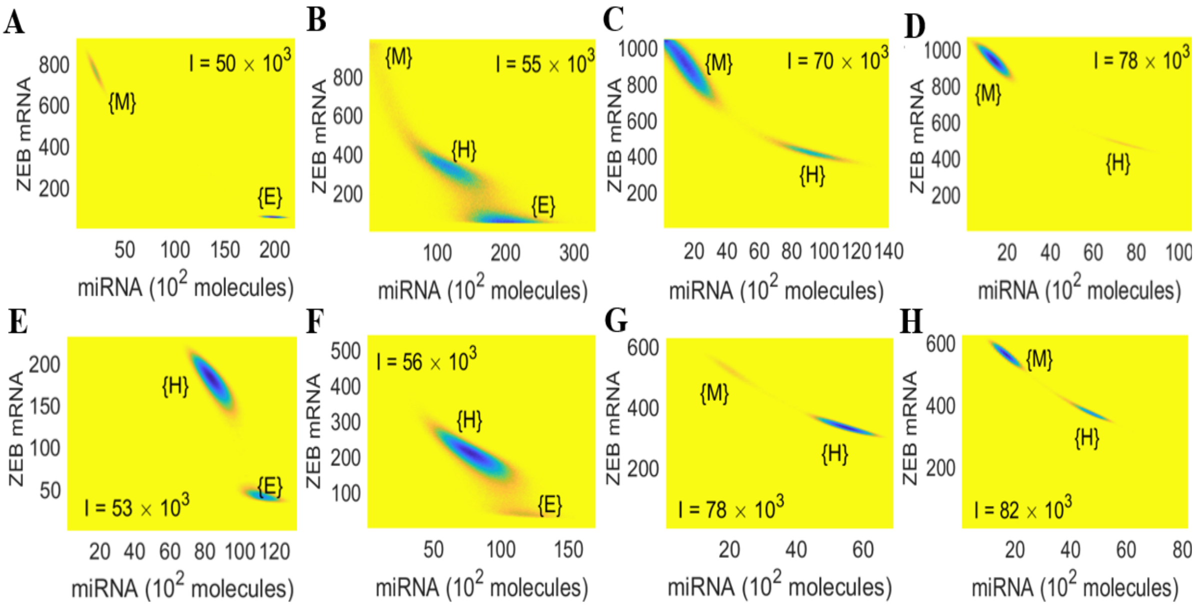


**Fig S4: Potential landscape A-D)** Potential landscape for core circuit at varying levels of external signal (I). **(E-H)** Potential landscape for the circuit including SLUG at varying levels of external signal (I).

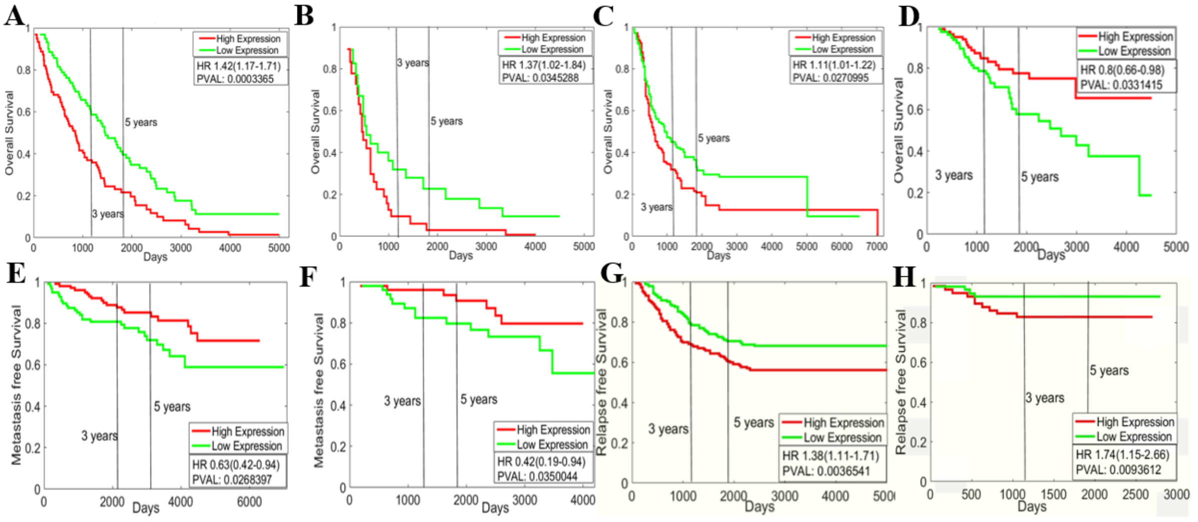


**Fig S5**: **Kaplan–Meier analysis for SLUG levels**. **A-D)** Overall survival of GSE26712 (ovarian cancer samples), GSE18520 (ovarian cancer sample), GSE13876 (ovarian cancer sample) and GSE3143 (breast cancer sample) respectively. **E-F)** Metastasis free survival of GSE11121 and GSE2990 (breast cancer samples) respectively. **G-H)** Relapse free survival of GSE2034 and GSE19615 (breast cancer samples) respectively. HR denotes Hazard ratio, PVAL denotes p-value.
